# Patterns of change in nucleotide diversity over gene length

**DOI:** 10.1101/2023.07.13.548940

**Authors:** Farhan Ali

## Abstract

Nucleotide diversity at a site is influenced by the relative strengths of neutral and selective population genetic processes. Therefore, attempts to identify sites under positive selection require an understanding of the expected diversity in its absence. The nucleotide diversity of a gene was previously found to correlate with its length. In this work, I measure nucleotide diversity at synonymous sites and uncover a pattern of low diversity towards the translation initiation site (TIS) of a gene. The degree of reduction in diversity at the TIS and the length of this region of reduced diversity can be quantified as “Effect Size” and “Effect Length” respectively, using parameters of an asymptotic regression model. Estimates of Effect Length across bacteria covaried with recombination rates as well as with a multitude of fast-growth adaptations such as the avoidance of mRNA secondary structure around TIS, the number of rRNAs, and relative codon usage of ribosomal genes. Thus, the dependence of nucleotide diversity on gene length is governed by a combination of selective and non-selective processes. These results have implications for the estimation of effective population size and relative mutation rates based on “silent-site” diversity, and for pN/pS-based prediction of genes under selection.

## I. INTRODUCTION

Populations in nature are known to be genetically polymorphic. The extent of this within-species polymorphism varies dramatically among loci across a genome [1]–[5]. As a result, there is considerable interest in identifying targets of positive natural selection based on their diversity relative to a set of known or assumed neutral loci [6]–[9]. Naturally, this endeavor would benefit from a better understanding of the processes that govern background genetic variation.

In our study on the variability of transcription factors and their target genes in *E. coli*, we found nucleotide diversity of genes to be positively correlated to their length and this effect was stronger for synonymous sites in comparison to non-synonymous sites [10]. Nucleotide diversity at a locus is calculated as the average pairwise difference per site among sequences in a population and so, is not expected to show any correlation with gene length. Since nucleotide diversity is now the fundamental measure of variation in population genetics [11, 12], and provides a basis for common tests of neutral evolution [13, 14], any factor that shapes its distribution within or across species must be clearly understood.

Variation in synonymous diversity across genes could be a reflection of varying degrees of selection on synonymous codons. Selection on codon usage has been observed to be correlated to gene length in *Escherichia coli* [15] and *Drosophila* [16]. Independently, a pattern of low synonymous substitution rates at both ends of a gene has been observed in many organisms [17]– 20], which is likely due to selection against mRNA secondary structure around ribosome-binding sites and initiation codons for efficient translation initiation [21]. Consequently, this is believed to operate within the first 50 bases of the translation initiation site (TIS) [18], and thus, would seem unlikely to be an explanation for the observed correlation of synonymous diversity with gene length. However, it could be that the region over which this selection operates extends beyond the first 50 sites and varies in length across species. We can resolve this issue by quantifying these patterns of synonymous polymorphism in a more systematic manner than attempted previously. More generally, the selection acting on translational efficiency in a bacterial species is expected to improve its growth rate. If this selection is indeed the primary force that drives the correlation between synonymous diversity and gene length, then we can expect these patterns to be more evident in species with higher growth rates, irrespective of the exact mechanisms.

In this paper, I first develop a method to quantify the average pattern of synonymous diversity over the length of *E. coli* genes. Then, I apply this method to multiple bacterial species in order to study how the strength of diversity-length correlation varies across species. After controlling for statistical correlates, I test the effect of recombination rates and several metrics of improved translation on the distribution of observed patterns of synonymous polymorphism across species.

## II. Methods

### Selection of strains

Genome assemblies were acquired from NCBI RefSeq, last accessed on Sep. 14, 2020. From the list of bacterial assemblies present on NCBI, the following were excluded: those missing from RefSeq, those that are miss-classified or uncultured, those without species name, or with the term “Candidatus” in place of a genus name, or “bacterium” in place of a species name. Out of the remaining species in the list, the ones with at least 30 chromosomal or higher level assemblies were selected, giving 75 species with a total of 9230 assemblies. For 24 species with more than 100 assemblies, 100 were randomly selected to reduce the overall computational burden, leaving a final set of 5272 genomes spread across 75 bacterial species.

For each species, genomes were further filtered to exclude highly similar ones. This was done based on the degree of overlap in their sets of chromosomal non-redundant protein ids. There are two sources of variation among these sets, *viz*., a) changes in the amino acid sequence of homologous proteins, and b) changes in the accessory genome. The pairwise dissimilarity between sets was measured with Jaccard distance, *i*.*e*., 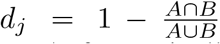 for protein sets corresponding to genomes A & B. Finally, a set of strains were selected for each species given a pairwise distance matrix such that they differed by at least 1 % in their set of protein ids *i*.*e*., *d*_*j*_ *≥* 0.01. This step brought down the total number of assemblies to 4939.

### Computing nucleotide diversity

Ortholog groups across selected strains of a species were identified using SonicParanoid (v1.3.8) in its fast mode with default parameters [22]. Single-copy orthologs, present in at least 75 % of the analyzed genomes, were selected for further analysis. Amino-acid sequences of orthologs were aligned using Clustal Omega (v1.2.4) [23] and converted to codon alignments using PAL2NAL (v14) [24]. Sites with over 70 % gaps were excluded. *N*-fold degeneracy of each site was determined using a consensus sequence derived from the alignment with a minimum consensus character threshold of 60 %. For each site, nucleotide diversity (*π*) was calculated as follows: 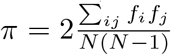, where the summation is over all possible pairs of 4 bases and *f*_*i*_ signifies their counts among *N* sequences.

### Effect length estimation

Diversity profiles over sites were quantified with respect to their effect size and length *i*.*e*., the magnitude and extent of the reduction in mean diversity observed near the start of a gene. Effect size is defined as the log-2 fold decrease in mean diversity at the start compared to the maximum mean diversity over the first 500 sites. Effect length is defined as the site at which the mean diversity attains the midpoint of its range. An asymptotic regression model (negative exponential) of the following form was used to fit observed mean diversities over 4-fold degenerate sites.

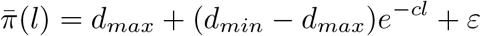

where 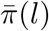 is the observed mean diversity at a site *l* bases downstream of the translation start site (index 0). *d*_*max*_ is the maximum mean diversity attainable as *l* goes to +, *d*_*min*_ is the minimum estimated mean diversity at *l* = 0, and *c* governs the relative rate of increase in diversity with distance from gene start. The error (*ε*) in estimating the site-specific mean is assumed to be additive, and independent with variance *σ*^2^*/w* [25]. Weights (*w*) are set to the number of genes for each site to take into account fluctuations in the mean due to varying sample sizes. Regression was performed using R’s nonlinear least squares function `nls` with its Gauss-Newton algorithm. Initial estimates of model parameters were obtained using R’s `SSasymp`. Effect size *S*_*e*_ and length *L*_*e*_ were calculated using estimates of *c, d*_*min*_ and *d*_*max*_, as follows:

The standard errors of these estimates can be derived from the standard errors of model parameters using the Delta method as follows:

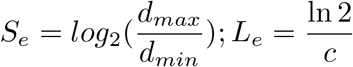

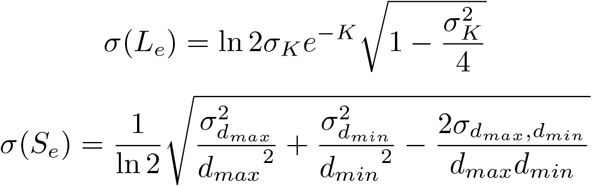

where *σ* stands for standard error and *K* = ln *c* is used since `SSasymp` fits the above model using ln *c*.

### Positional probabilities for RNA-secondary structure

Vienna RNA package (v2.4.16) was used to predict RNA secondary structures [26]. The probability of each site remaining unpaired (open) in an RNA secondary structure was computed for every gene in a genome over a region from 100 bp upstream to 200 bp downstream (−100:+200) of the translation start site. Partition functions were used to get the base-pairing probabilities of each site in a thermodynamic ensemble of secondary structures. Probabilities were converted to z-scores following Molina’s method [18], with the flanking regions marked as -90 to -51 and +151 to +190. A z-statistic profile was generated for one reference genome from each species. Effect length corresponding to selection on mRNA secondary structure *L*_*s*_ was defined as the first site +5 onwards at which the z-score drops to zero; effect size *S*_*s*_ was measured as the maximum z-score over the same region.

### Estimation of recombination rate

Recombination rate for each species was estimated from codon alignments, using the Mcorr program [27]. Alignments of individual genes were first concatenated in a single alignment in XMFA format, which was submitted as input to the program. Mcorr takes a coalescent approach to estimate the mutational (*θ*_*p*_ = 2*N*_*e*_*μ*) and recombinational divergence (*ϕ*_*p*_ = 2*N*_*e*_*γ*) of a bacterial population given a sample of genomes, by fitting an analytical model to substitution correlation profile among pairs of synonymous sites. The rate of recombination relative to mutation rate (r/m) was estimated as *ϕ*_*p*_*/θ*_*p*_. Estimates were log-transformed (Box-Cox’s *λ* = 0) for regression analysis.

### Estimation of codon-usage bias and minimum doubling times

Codon-usage bias (CUB) was estimated using gRodon package in R [28]. gRodon uses CUB of highly expressed genes, and other associated variables, to predict minimum doubling times based on a regression model trained on growth rate data from Vieira-Silva & Rocha 2010, and Madin *et al* 2020 [29, 30]. For each genome, its multi-FASTA file of coding sequences was used as input to gRodon. First, ribosomal genes were used as the set of highly expressed genes to calculate CUB, which was then used to predict minimum doubling times *DT*_*pred*_. Estimates of *CUB*_*HE*_ and *DT*_*pred*_ were averaged over all genomes for each species. A squared-root transformation was applied to *CUB*_*HE*_ (Box-Cox’s *λ* = 0.5) prior to regression analysis, and *DT*_*pred*_ was log-transformed (Box-Cox’s *λ* = *−*0.1).

### Phylogenetic reconstruction

UBCG program was used to reconstruct bacterial phylogeny based on concatenated amino acid alignments of 81 universal bacterial markers [31]. Only one reference genome assembly was used for each bacterial species. In the absence of a reference assembly, as defined in the NCBI RefSeq database, a representative genome was used. For the two species (*Planctomycetes* and *Wolbachia*) that lacked both a reference and a representative genome, the genome with the number of genes closest to the mean was selected. Contigs identified as plasmid by the NCBI annotation pipeline were removed from these reference sequences. Positions with more than 50 % gaps were removed from the alignment. A maximum likelihood phylogeny was estimated using the RAxML program (v8.2.12), with a Jones-Taylor-Thornton (JTT) model of sequence evolution, along with CAT approximation [32]. UBCG provides branch support in terms of a gene support index (GSI), *i*.*e*., the number of gene trees supporting a bipartition in the species tree. The default threshold of a minimum of 95 % similarity was used to decide whether a branch is supported by a gene tree. The tree was rooted in between Terrabacteria and Gracilicutes, following a recently published rooted phylogeny of all bacteria [33].

### Phylogenetic comparative analysis

A phylogenetic generalized least squares approach (phylo-GLS) was used to test for relationships among variables across species while accounting for their phylogenetic correlation. The strength of the phylogenetic signal on the distribution of a genetic trait or residuals of a regression model can be quantified using Pagel’s *λ* [34]. Pagel’s *λ* is a multiplier of the off-diagonal entries of the phylogenetic covariance matrix wherein a value of 0 signifies independent evolution and a value of 1 signifies the evolution in complete accordance with Brownian motion over shared ancestry [35]. `gls` function from R package “nlme” was used to perform generalized least squares, and `corPagel` function from R package “ape” was used to calculate the required correlation structure based on Pagel’s *λ* [36].

### Statistical analysis

Variables were transformed, wherever necessary, using Box-Cox transformation to reduce skewness and approximate normality. Variance Inflation Factor (VIF) was used to check and control for strong collinearity among predictors. VIF for a variable is calculated as 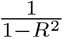 of a linear regression in which it appears as a response variable dependent on the rest of the predictors. Regression was performed on scaled variables (using mean and standard deviation) to make coefficients comparable. All statistical analyses were performed using R (v4.1.2).

## III. Results

### A. Diversity-length correlation reflects reduced polymorphism toward gene starts

We have seen previously that nucleotide diversity is positively correlated with gene length in *Escherichia coli* [10]. The strength of this correlation was stronger for synonymous sites, suggesting that the process driving this length-dependence was directly acting on the nucleotide sequence. In this study, I estimate diversity from 4-fold degenerate sites to eliminate the effect of any selection acting on amino-acid sequences. For a collection of 96 *E. coli* genomes, selected as described in Methods, the positive correlation between gene length and silent-site diversity was evident, with a distinctly non-linear, saturating trend **Figure 1**. This could result from an underlying pattern of site-to-site variation in nucleotide diversity such that shorter genes have a smaller proportion of high-diversity sites. Such patterns of reduced variation near translation initiation sites (TIS) have been observed previously [17]– 20]. The primary question here is whether these patterns can explain the observed diversity-length correlation. Since the patterns over individual genes are highly variable [**SI Figure 1**], I instead focus on the trend of average nucleotide diversity over sites, quantify the magnitude and reach of this effect, and compare patterns across species to understand the basis of the observed correlation between gene length and nucleotide diversity.

**Fig. 1:**
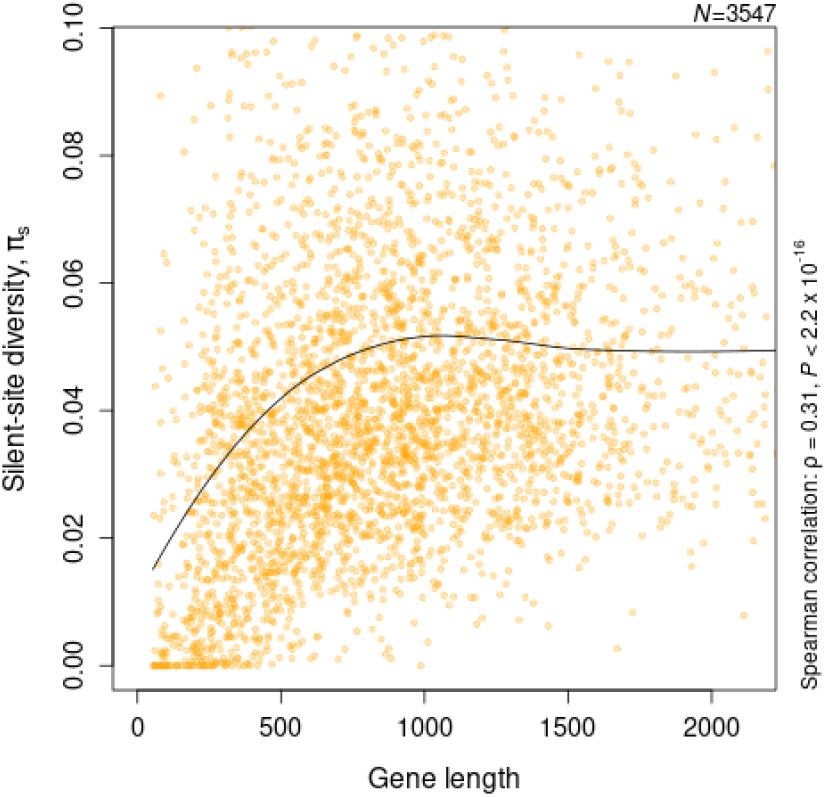
Correlation between nucleotide diversity and gene length. Nucleotide diversity was estimated from silent sites (4-fold degenerate) of 3547 genes present in at least 75 % of the 96 *E. coli* genomes. X- and Y-axis are limited to their respective upper 95th percentile for visualization. The black line shows the LOESS curve with a span of 0.75 and highlights the broad trend of change in synonymous diversity with gene length.

As an asymptotic trend in the nucleotide diversity of a site averaged over genes is apparent from **Figure 2A**, a natural choice of fit for diversity-site profiles would be the asymptotic regression (ASR) model. Such a negative exponential equation can be used to quantify the magnitude and extent of this effect on diversity, which can be compared across samples of strains or species. Here onward, “Effect length *L*_*e*_” is defined as the number of silent sites from the translation starts at which the estimated average diversity is halfway to saturation, and “Effect size *S*_*e*_” is defined as the log-2 fold difference between the maximum and minimum estimated mean diversity. Effect length in *E. coli* was estimated as equal to 76 [**Figure 2A**]. The point of saturation can be conservatively approximated as 4*L*_*e*_ = 304. The estimated Effect Size of 2.2 captures the nearly 5-fold decrease in diversity at the start site relative to its expected value at saturation.

**Fig. 2:**
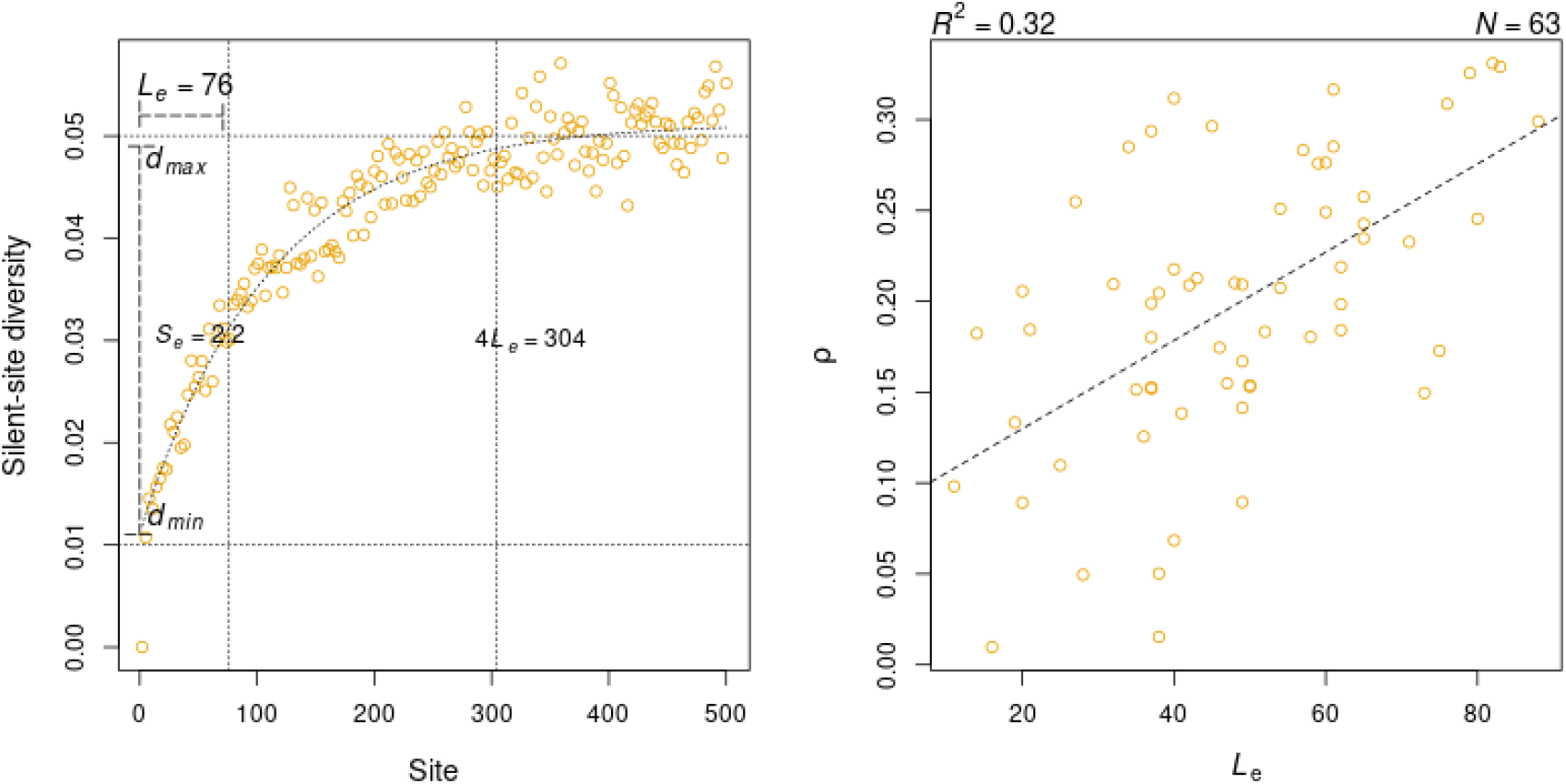
Patterns of reduced diversity toward the start of bacterial genes. **A)** Asymptotic regression fit to mean nucleotide diversity of 4-fold degenerate sites averaged over *E. coli* genes. *L*_*e*_ and *S*_*e*_ mark the estimates of Effect Length and Size respectively, quantifying the magnitude and extent of the observed reduction respectively. **B)** Linear regression of the strength of correlation between gene length and silent-site diversity on estimates of *L*_*e*_ for 63 bacterial species with *S*_*e*_ *>* 0 and *ρ >* 0.

Using 75 species with at least 30 genomes, the ASR model described above could be fit to the mean nucleotide diversity profile of 69 species, 65 of which had a positive Effect Size. Thus, over 85 % of the analyzed species showed clear signs of reduced polymorphism towards the 5’-end of genes. The estimates of Effect Length ranged from 11 in *Streptococcus thermophilus* to 88 in *Acinetobacter baumannii*. The variability in Effect Length estimates across species was greater than that of the Effect Size, both in terms of their observed coefficients of variation, 0.39 v/s 0.32, and as the ratios of observed standard deviation to the corresponding standard error in *E. coli*, 4.72 v/s 3.94. Estimates of *L*_*e*_ and *S*_*e*_ across species were correlated to each other (Spearman correlation, *ρ* = 0.46), as well as to the strength of the correlation between gene length and nucleotide diversity (*ρ*_*Le*_ = 0.54, *ρ*_*Se*_ = 0.45). Multiple linear regression of this correlation coefficient (*ρ*_*l*_) on *L*_*e*_ and *S*_*e*_ found the effect of *L*_*e*_ to be much stronger than that of *S*_*e*_ (*P*_*Le*_ = 0.0004, *P*_*Se*_ = 0.0462). A simple linear regression with *L*_*e*_ alone explained nearly one-third of the variation in the degree of gene-length dependence of nucleotide diversity [**Figure 2B**]. The rest of this study is focused on understanding the factors that shape the distribution of Effect Length across species.

### B. Effect length is shaped by selection on mRNA secondary structure and the rate of recombination

The observed distribution of Effect Lengths across species could be biased by confounding variables. Before proceeding with understanding the effect of biological factors, I tested for the correlation of Effect Length with potential statistical confounders, such as the number of strains, number of genes, average gene length, mean nucleotide diversity, and GC content [**SI Text 1**]. Only the number of strains had a significant correlation with Effect Length and was controlled for in further analyses [**SI Table 1**].

**TABLE I:** Spearman correlation among Effect Length and its covariates.

| | $S_{ss}$ | $\ln r/m$ | $\sqrt{N_{rrn}}$ | $\sqrt{CUB_{HE}}$ |
| --- | --- | --- | --- | --- |
| $\ln r/m$ | 0.04 | | | |
| $\sqrt{N_{rrn}}$ | 0.25 | -0.04 | | |
| $\sqrt{CUB_{HE}}$ | 0.38 | -0.27 | 0.58 | |
| $L_e$ | 0.26 | -0.35 | 0.47 | 0.51 |

The most common explanation for reduced polymorphism at 5’-ends of protein-coding genes seems to be the selection to avoid mRNA secondary structure formation around TIS [17, 18, 37]. However, this selection is believed to operate within the first 50 bp of CDS, and consequently, attempts to quantify the magnitude and extent of this effect have focused only on this limited region. To test if the selection of mRNA secondary structure can explain the variation in *L*_*e*_ across species, I used RNA secondary structure prediction over a larger region *i*.*e*. 100 bases upstream to 200 bases downstream of TIS, and estimated the average secondary-structure free length (*L*_*ss*_) for each species. More specifically, I calculated the probability of each base being unpaired across an ensemble of possible RNA secondary structures over the specified region of every gene in a genome. I converted raw probabilities to Z-scores following Molina’s approach [18], such that the Z-score measures the tendency of a base to be unpaired relative to the bases at the ends of the segment. I defined Effect Length (*L*_*ss*_) and Effect Size (*S*_*ss*_) in this context as the first site at which Z-score drops to 0 and as the maximum Z-score respectively. *L*_*ss*_ for 75 species ranged from 23 to 114, with limited dispersion, as indicated by its inter-quartile range (IQR = *Q*_3_ *− Q*_1_) of 9, relative to that of *L*_*e*_ with IQR = 24 [**Figure 3B**]. *L*_*e*_ has a negligible correlation with *L*_*ss*_ (*ρ* = *−* 0.08, *P* = 0.51, *N* = 64, Spearman correlation) but a significant correlation with *S*_*ss*_ (*ρ* = 0.27, *P* = 0.03). Even when accounting for the number of strains in a multiple regression model, the effect of *S*_*ss*_ on *L*_*e*_ remains significant (*P* (*S*_*ss*_) = 0.0187, *P* (*N*_*strains*_) = 0.0048). Hence, while the length over which the selection to avoid mRNA secondary structure operates is relatively more uniform, patterns of low diversity near TIS do exhibit a weak dependence on the strength of this effect across species.

**Fig. 3:**
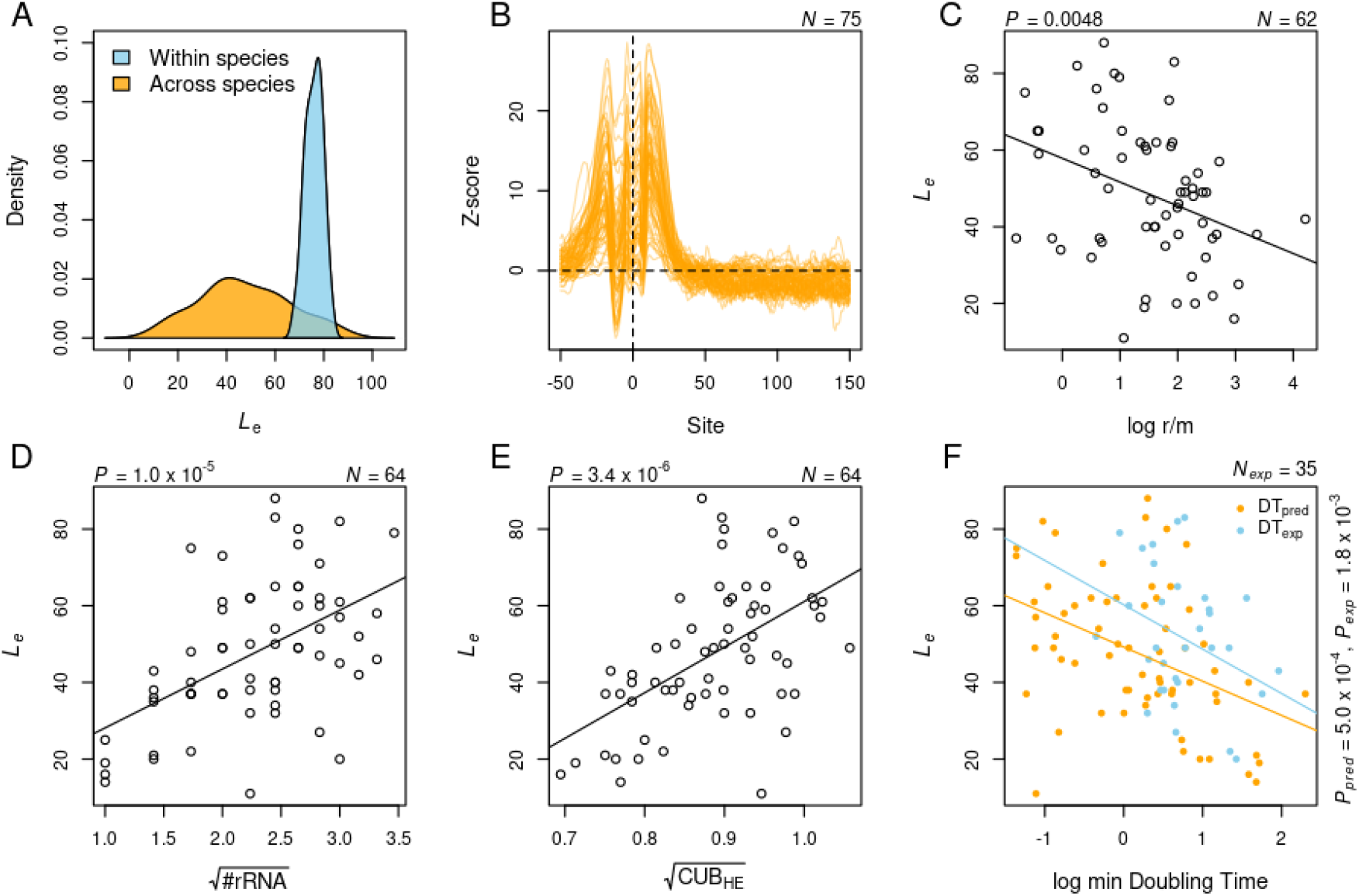
Effect Length and its predictors across species. **A)** Distribution of Effect Length within and across species. Within-species distribution is represented by a random sample, of the same size as the number of species, drawn from a Normal distribution with mean and standard deviation equal to the estimate of Effect Length in *E. coli* and its standard error respectively. Across-species distribution is more dispersed with its lower range of values well beyond the distribution of Effect Length in *E. coli*. **B)** Selection to avoid mRNA secondary structure around TlSS. Z-score profiles for the probability of a base being unpaired in an mRNA secondary structure were drawn for 75 species. Z-scores for different species show considerable variation in their maximum values but attain baseline within a narrow range of positions. **C)** Variation in Effect Length can be partially explained by species-specific recombination rate. The recombination rate was measured using Mcorr as a log-ratio of the rate of recombination to mutation **D) & E)**. Effect Length was strongly correlated with rRNA count and relative codon usage bias of highly expressed genes respectively. Both predictors were squared-root transformed based on their Box-Cox *λ*. **F)** Effect Length is negatively correlated with predicted as well as experimentally observed minimum doubling times. Doubling times are in hours. Natural logarithm of doubling times was taken for appropriate Box-Cox transformation. All mentioned p-values are for simple linear regression of Effect Length on a given predictor.

Even if the strength of selection was uniform, Effect Length could still differ among species due to the varying extent of linkage disequilibrium. If recombination is infrequent, then even the neutral sites linked to loci under purifying selection will have reduced polymorphism [38]. I tested this idea by estimating the ratios of recombination-to-mutation rate per nucleotide site using Mcorr [Methods] [27]. Mcorr uses a coalescent-based approach to estimate the recombinational and mutational divergence of a bacterial population given a sample. Out of 64 species with Effect length, r/m of 2 species had extreme values *viz*., *Bordetella parapertussis* (*<* 10^*−*3^) and *Wolbachia* (*>* 10^3^), leaving 62 species. Linear regression along with the number of strains identified a significantly negative effect of r/m on *L*_*e*_ (*P* = 0.0097) [**Figure 3C**]. The model had a strong phylogenetic signal (Pagel’s *λ* = 0.7), even accounting for which didn’t eliminate the effect of recombination rate on *L*_*e*_ (*P* = 0.0152). Therefore, low levels of recombination extend the regions of reduced diversity around TlSS beyond the short length over which the selection to avoid mRNA secondary structure operates.

### C. Effect length reflects growth phenotype of a species

The avoidance of the mRNA secondary structure around the translation start site, as discussed above, is a mechanism to improve translation initiation rates through greater accessibility of ribosomes to their binding sites [39]. All else being equal, a more efficient translation should lead to faster growth of the species. Since a greater magnitude of this selection leads to a longer Effect Length, one might expect Effect Length to correlate with improved growth phenotypes. However, the growth characteristics of the majority of bacteria are unknown as they remain unculturable, and even for those that can be cultured, their growth potential might not be realized in laboratory conditions. In the absence of experimentally verified growth rates for many species, I used genome-based predictions of minimum doubling times to test this idea. More specifically, I used the gRodon program that employs relative codon usage bias of highly expressed genes (CUB-HE) and additional metrics to predict the minimum doubling time for a given genome [28]. Additionally, I used the average rRNA count (*N*_*rrn*_)for each species as the number of rRNAs is known to correlate with faster growth [29]. All of the 4 variables *viz*., CUB-HE, *N*_*rrn*_, *L*_*e*_, and predicted minimum doubling times (*DT*_*pred*_) were correlated to each other [**Figure 3D-F, Table I**]. *L*_*e*_ was more strongly correlated with CUB-HE and *N*_*rrn*_ than with *DT*_*pred*_ itself. Considering collinearity among CUB-HE, *N*_*rrn*_ and *DT*_*pred*_, Variance Inflation Factor (VIF) of CUB-HE and *DT*_*pred*_ was comparable to each other and greater than that of *N*_*rrn*_ [**SI Table 2**]. Since *DT*_*pred*_ is additionally a composite measure based on other genome characteristics, I chose to include CUB-HE in the following regression analysis instead of *DT*_*pred*_. Multiple linear regression of *L*_*e*_ on *N*_*rrn*_ and CUB-HE, along with *N*_*strains*_ found all three predictors to be relevant [**SI Table 2**]. The residuals of this model showed a weak phylogenetic signal (Pagel’s *λ* = 0.1(*−*0.19, 0.4 : 95%*CI*)), even accounting for which didn’t change the above results.

**TABLE II:** Regression models of Effect Length on different subsets of predictors. The models with minimum test Root Mean Squared Error (RMSE) in each category of the number of predictors have been listed. Plus(+) signs mark the variables corresponding to each model. The last column lists the Mean Absolute Error of each model (MAE).

| | $S_{ss}$ | $\ln r/m$ | $\sqrt{N_{rrn}}$ | $\sqrt{CUB_{HE}}$ | $N_{strain}$ | RMSE | $R^2$ | MAE |
| --- | --- | --- | --- | --- | --- | --- | --- | --- |
| 1 | - | - | + | - | - | 16.44 | 0.31 | 13.26 |
| 2 | - | + | + | - | - | 16.12 | 0.32 | 12.90 |
| 3 | - | + | + | - | + | 15.88 | 0.33 | 12.71 |
| 4 | - | + | + | + | + | 14.50 | 0.43 | 11.56 |
| 5 | + | + | + | + | + | 14.38 | 0.43 | 11.37 |

The application of genomic attributes to test for a relationship between Effect Length and maximum growth rate is not entirely satisfactory since it is possible that the Effect Length is associated with other aspects of these genomic traits that are unrelated to their effect on growth rates. Any relationship between Effect Length and growth rates would be far more convincing if corroborated with experimental data. Minimum doubling times determined through cell cultures are indeed available for at least half of the species used in this study. However, one of the main issues in using experimental growth rates is the variation caused due to different laboratory growth conditions, in particular, temperature. In the most recent collection of experimental rates, a correction for differences in temperature was applied by normalizing all growth rates to 20^*?*^C assuming *Q*_10_ = 2.5. Such normalized growth rates were available for 31 of the species used in this study. For another 4 species, growth rates were taken from a previous collection [29] and normalized for temperature using the same approach as above. Finally, with these experimentally determined, minimum doubling times (h) of 35 species, I tested for a relationship between Effect Length and maximum growth rates. Effect Length was indeed correlated with minimum doubling times (*ρ* = *−* 0.5, *P* = 0.004, Spearman correlation. *R*^2^ = 0.24), suggesting that Effect Length might reflect the strength of selection on the growth characteristics of an organism.

### D. Fast growth adaptations coupled with recombination rates shape the distribution of Effect Length across species

Having studied the association of individual factors with Effect Length and found a set of key correlates, the final step was to identify the most effective predictors and their independent effects. Multiple linear regression of *L*_*e*_ on *S*_*ss*_, *r/m, N*_*rrn*_, *CUB*_*HE*_ and *N*_*strains*_ using 62 species identified rRNA count and recombination rate as the top two significant predictors followed by the number of strains. I compared the full model against simpler models using a backward elimination procedure with a 10-fold cross-validation repeated 100 times. The full model turned out to be the best model as judged by its minimum prediction RMSE (*RMSE* = 14.38, *R*^2^ = 0.43), with rRNA count and recombination rate as the two most effective predictors [**Table II**].

The phylogenetic signal in this model was weak enough to be ignored in favor of a simpler model (Pagel’s *λ* = 0.1 (−0.31:0.52, 95% CI)). However, this lack of phylogenetic signal is expected with a large number of parameters and few observations, and thus cannot be taken as evidence of phylogenetic independence of the distribution of Effect Length as governed by the above factors. To test for true independence, I split the dataset by the two major phylogenetic groups *i*.*e*. Terrabacteria (24) and Gracilicutes (38). Gram-positive and Gram-negative bacteria belong to the groups Terrabacteria and Gracilicutes respectively. The full models trained on one group and tested on the other performed poorly (*R*^2^ ≊ 0.07), which highlights the phylogenetic dependence of these relationships. While these individual models are useful for demonstrating the phylogenetic dependence of Effect Length on its covariates, they lack the power to reveal all relevant relationships in this limited dataset. In the absence of extensive population genomic data presently, all of the above predictors should be considered relevant to the full distribution of Effect Length across species. Therefore, a pronounced lack of polymorphism at silent sites toward 5’-ends of protein-coding genes is a reflection of the combined action of fast-growth adaptations and recombination rate.

## IV. Discussion

In this study, I sought to identify a basis for a positive correlation between nucleotide diversity and gene length observed previously [10]. The pattern of reduced polymorphism toward the translation start site explains a considerable proportion of this gene-length dependence. Fitting an asymptotic regression model to the observed pattern of mean diversity over sites enabled quantification of the size (*S*_*e*_) and length (*L*_*e*_) of this effect across species. With a phylogenetic comparative approach, I found the distribution of Effect Length to be shaped by a combination of selective and non-selective processes.

Effect Lengths of different bacteria showed a positive correlation with their maximal growth rates. Species with smaller doubling times are under stronger selection for optimizing translation rates, and Effect Length could be a reflection of the strength of this selection. This view is supported by correlations of Effect Length with other fast-growth adaptations, such as rRNA copy number, and codon-usage bias of highly expressed genes. Alternatively, the strength of selection acting on the starting region of genes could be the same across species but the countering effect of random genetic drift is weaker in fast-growing bacteria. Indeed, Effective population size (*N*_*e*_) was found to correlate positively with growth rate across bacterial species [40]. When *N*_*e*_ is higher, variants that increase the stability of RNA secondary structures around translation start sites can be more effectively removed from the population, resulting in a clear pattern of reduced polymorphism in the starting region of genes, and reflected in the higher estimates of *L*_*e*_. Eyre-walker found synonymous codon bias in *E. coli* to positively correlate with gene length among genes expressed at comparable levels and argued that it reflected selection to avoid missense errors in longer proteins due to their higher cost of production [15]. While this could certainly be true for the small set of proteins used in his study, especially ribosomal proteins, it is expected to lead to a negative correlation between silent-site diversity and gene length on a genome-wide scale, contrary to our observations. The counter-acting selection towards the 5’-end of genes could explain this pattern, such that shorter genes have a lower bias due to conflicting forms of selection, but he assumed it to be relevant for the first 50 bases. The selection to avoid mRNA secondary structure around TIS does indeed appear to be limited to this range [18, 37]. However, nucleotide diversity of linked neutral sites further downstream in the gene would also be reduced due to this purifying selection [38]. Indeed, the wide range of variation in recombination rates across bacteria [41] seems to have shaped the Effect Length distribution, such that species with low recombination frequency are more likely to show higher Effect Length.

Estimates of synonymous diversity have been used to study mutation rate variation across genes after controlling for many suspected factors [2]. However, gene length was not controlled for in that study and shows a significant correlation with the estimates of synonymous diversity, even after correcting for other correlates (*ρ* = 0.15, *P* = 1.64 *×* 10^−14^, Spearman rank correlation). Admittedly, there’s still substantial unexplained variation in the synonymous diversity of genes which highlights the need for continued efforts in understanding selective and neutral processes that shape nucleotide composition and patterns of diversity [42]– 45].

To study the efficacy of natural selection in driving any genetic trait to fixation, evolutionary biologists need to know the effective population size. Under mutation-drift equilibrium, the nucleotide heterozygosity of a bacterial population is expected to approximate 2*N*_*e*_*μ*, where *μ* is the per-base pair mutation rate per generation [46]. Thus, given experimental measures of *μ, N*_*e*_ is commonly estimated by equating the above expression with the nucleotide diversity of 4-fold degenerate sites assuming their near-neutrality [47, 48]. However, as quantified in this study by the effect size, estimates of *N*_*e*_ at the start of bacterial genes can be nearly 5-fold lower than the average [49]. Given that the extent of this effect varies substantially across species, these regions of reduced polymorphism should be separately identified and removed before effective population sizes of different species are compared.

One final issue with ignoring the non-neutrality of synonymous diversities and their gene-length dependence concerns the detection of genes under positive selection. Common tests of the strength of selection acting on a protein-coding sequence involve the ratio of the rate of non-synonymous substitutions to that of synonymous substitutions [50]– 52]. Often, the ratio of non-synonymous to synonymous diversity is used instead of substitution rates [53]– 55], or the method is used for intraspecific comparisons [56, 57], both of which can be misleading in their own [58, 59]. However, the pattern of reduced variation toward gene starts studied here using synonymous diversity is also visible when synonymous substitution rates are used instead [17]. Therefore, the extent of purifying selection on synonymous sites seen in this study could lead to an overestimation of the prevalence of positive selection [60]. Increasing recognition of potentially unknown sources and consequences of even weak synonymous selection continues to advance our ability to identify genuine cases of molecular adaptation [61].

## V. Data Availability

The source codes, plotting scripts, metadata, and analysis outputs of this study can be accessed from the GitHub repository A-Farhan/diversity length correlation.

## VI. Acknowledgements

This research was facilitated by the National Centre for Biological Sciences, Tata Institute of Fundamental Research, India, and by the Biodesign Centre for Mechanisms of Evolution, Arizona State University, USA. The author is currently supported through the Multidisciplinary University Research Initiative award W911NF-14-1-0411 from the US Army Research Office, National Institutes of Health award R35-GM122566-01, National Science Foundation award DBI-2119963, and Grant 735927 from the Moore-Simons Project, awarded to Michael Lynch. I thank Aswin Sai Narain Seshasayee and Michael Lynch for their helpful comments on an earlier version of this manuscript.

