## Supplemental Text for "Patterns of change in nucleotide diversity over gene length"

**SI Text 1: Statistical correlates of $L_e$**

The observed distribution of Effect Lengths across species could be biased by potential confounding variables involved in this analysis, such as the number of strains, number of genes, average gene length, and mean diversity. Additionally, as the frequency of 4-fold degenerate sites is governed by the GC content, such that high GC species will have a higher density of such sites on average within a given coding region, Effect Length could be correlated to GC content as well. Spearman correlation of these 5 variables *viz.*, the number of strains $N_{strains}$, the number of genes $N_{genes}$, average gene length $L_{gene}$, average gene diversity $\boldsymbol{\pi}_{gene}$, and GC content $P_{GC}$ with Effect Length was non-significant after multiple-testing correction (Holm-Bonferroni's method), except that for the number of strains and the number of genes [**SI Table 1**]. To be able to control for these correlations in further analyses, and to estimate the proportion of variance in the Effect Length across species explained by these variables alone, a multiple linear regression analysis among these confounders and the Effect Length was performed. First, the distributions of individual variables were analysed for their shapes and the presence of outliers. Nucleotide diversity was log-transformed (Box-Cox $\lambda \approx 0$). *Planctomycetes* bacterium was excluded, on account of its average gene length of 1565 that is 6 standard deviation (SD) greater than the mean of 974 across species. Then, collinearity among predictors was checked by calculating Variance Inflation Factor (VIF) for each predictor variable. VIF for a variable is calculated from a linear regression in which it appears as a response variable dependent on the rest of the predictors. It is calculated as $1/(1-R^2)$ from the corresponding model and is supposed to be high if other predictors can explain a large proportion of the variance in the target variable. GC content had the highest VIF of 2.3, followed by the number of genes (2.1) [**SI Table 1**]. Since GC content also had the weakest Spearman correlation with Effect Length, it was excluded from further analyses. Finally, linear regression of $L_e$ on $N_{strains}$, $N_{genes}$, $L_{gene}$, and $\boldsymbol{\pi}_{gene}$ using their scaled distributions (z-scores) revealed that at least one of these factors was significantly associated with the Effect Length ($F$-statistic = 4.843, $P$ = 0.001911). The linear model explained 20 % of the variation in $L_e$ ($R^2$ = 0.196) and the greatest effect was due to $N_{strains}$ and $N_{genes}$.

Effect length appears to be correlated to both the number of strains and the number of genes. However, these analyses assumed independence of observations which may not be true for many genetic traits. The strength of phylogenetic signal in the above model was quantified using Pagel's $\lambda$. For a Generalised Least-Squares (GLS) regression of $L_e$ on $N_{strains}$, $N_{genes}$, $L_{gene}$, and $\boldsymbol{\pi}_{gene}$, the maximum likelihood estimate of Pagel's $\lambda$ to be 0.48, significantly greater than 0 (Likelihood ratio test: $\boldsymbol{\Lambda}$ = 5.98, $P_{\chi 2}$(df=1) = 0.014). With this model, $N_{genes}$ was no longer significant [**SI Table 1**]. However, $N_{strains}$ still had a significant positive effect on $L_e$ ($P$ = 0.004) and accordingly, was controlled for in further analyses.

**SI Table 1: Statistical correlates of Effect Length.**

| | Spearman correlation | | | Collinearity | | Linear Regression | | phylo-GLS | |
|---|---|---|---|---|---|---|---|---|---|
| Variables | $\boldsymbol{\rho}$ | $P_{raw}$ | $P_{adjusted}$ | $R^2$ | VIF | $\boldsymbol{\beta}$ | P($\|\boldsymbol{\beta}\|$>0) | $\boldsymbol{\beta}$ | P($\|\boldsymbol{\beta}\|$>0) |
| $N_{strains}$ | 0.35 | 0.004 | 0.02 | -0.010 | 0.990 | 0.315 | 0.009 | 0.31 | 0.004 |
| $N_{genes}$ | 0.32 | 0.01 | 0.042 | 0.518 | 2.074 | 0.290 | 0.017 | 0.24 | 0.094 |
| $L_{gene}$ | -0.29 | 0.02 | 0.061 | 0.397 | 1.660 | -0.093 | 0.448 | -0.03 | 0.835 |
| $\boldsymbol{\pi}_{gene}$ | -0.27 | 0.031 | 0.061 | 0.135 | 1.156 | -0.144 | 0.234 | -0.16 | 0.156 |
| $P_{GC}$ | 0.03 | 0.838 | 0.838 | 0.574 | 2.348 | NA | NA | NA | NA |

**SI Table 2: Growth rate correlates of Effect Length.**

| | Spearman correlation | | | Collinearity | | Linear Regression | | phylo-GLS | |
|---|---|---|---|---|---|---|---|---|---|
| Variables | $\boldsymbol{\rho}$ | $P_{raw}$ | $P_{adjusted}$ | $R^2$ | VIF | $\boldsymbol{\beta}$ | P($\|\boldsymbol{\beta}\|$>0) | $\boldsymbol{\beta}$ | P($\|\boldsymbol{\beta}\|$>0) |
| $N_{rrn}$ | 0.50 | $2.3 \times 10^{-5}$ | $6.9 \times 10^{-5}$ | 0.39 | 1.63 | 0.254 | 0.0522 | 0.262 | 0.0552 |
| CUB-HE | 0.53 | $7.4 \times 10^{-6}$ | $2.9 \times 10^{-5}$ | 0.72 | 3.62 | 0.364 | 0.0061 | 0.356 | 0.0082 |
| $DT_{pred}$ | -0.43 | $4.5 \times 10^{-4}$ | $9.0 \times 10^{-4}$ | 0.71 | 3.42 | NA | NA | NA | NA |
| $N_{strains}$ | 0.35 | $5.0 \times 10^{-3}$ | $5.0 \times 10^{-3}$ | -0.02 | 0.98 | 0.253 | 0.0142 | 0.266 | 0.0088 |

**SI Figure 1: Patterns of nucleotide diversity over 4-fold degenerate sites**, for 25 randomly selected *E. coli* genes. Value for each site is an average over the next 120 bases.

**SI Figure 2: Maximum-likelihood phylogeny of 75 bacterial species used in this study.** The tree was rooted in between Terrabacteria and Gracilicutes. Bootstrap values for a branch represent the number of gene trees out of 81 universal markers supporting the branch in the species tree.
