## Supplementary figures and images for "Patterns of change in nucleotide diversity over gene length"

### SI Figure 1

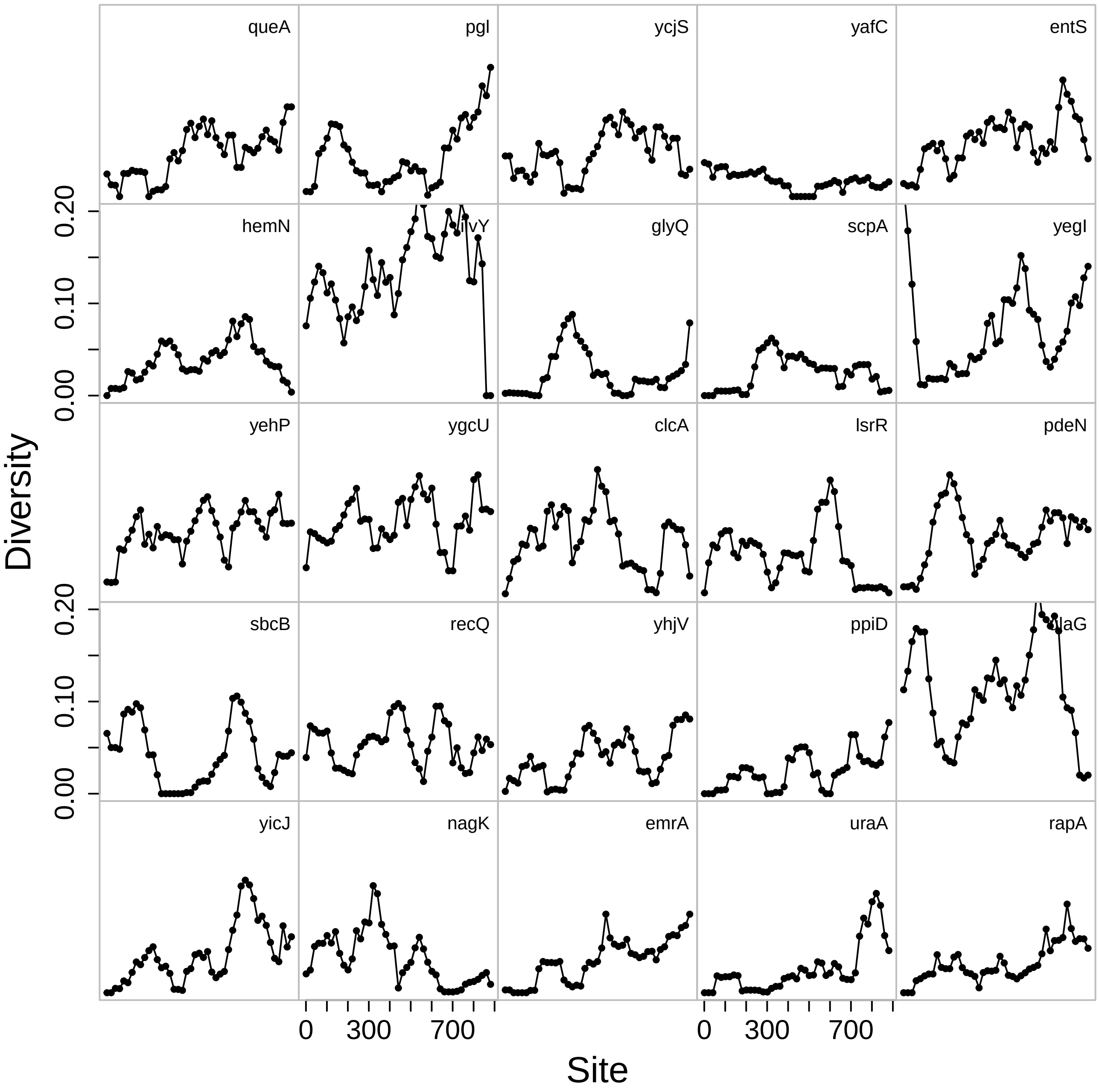

### SI Figure 2

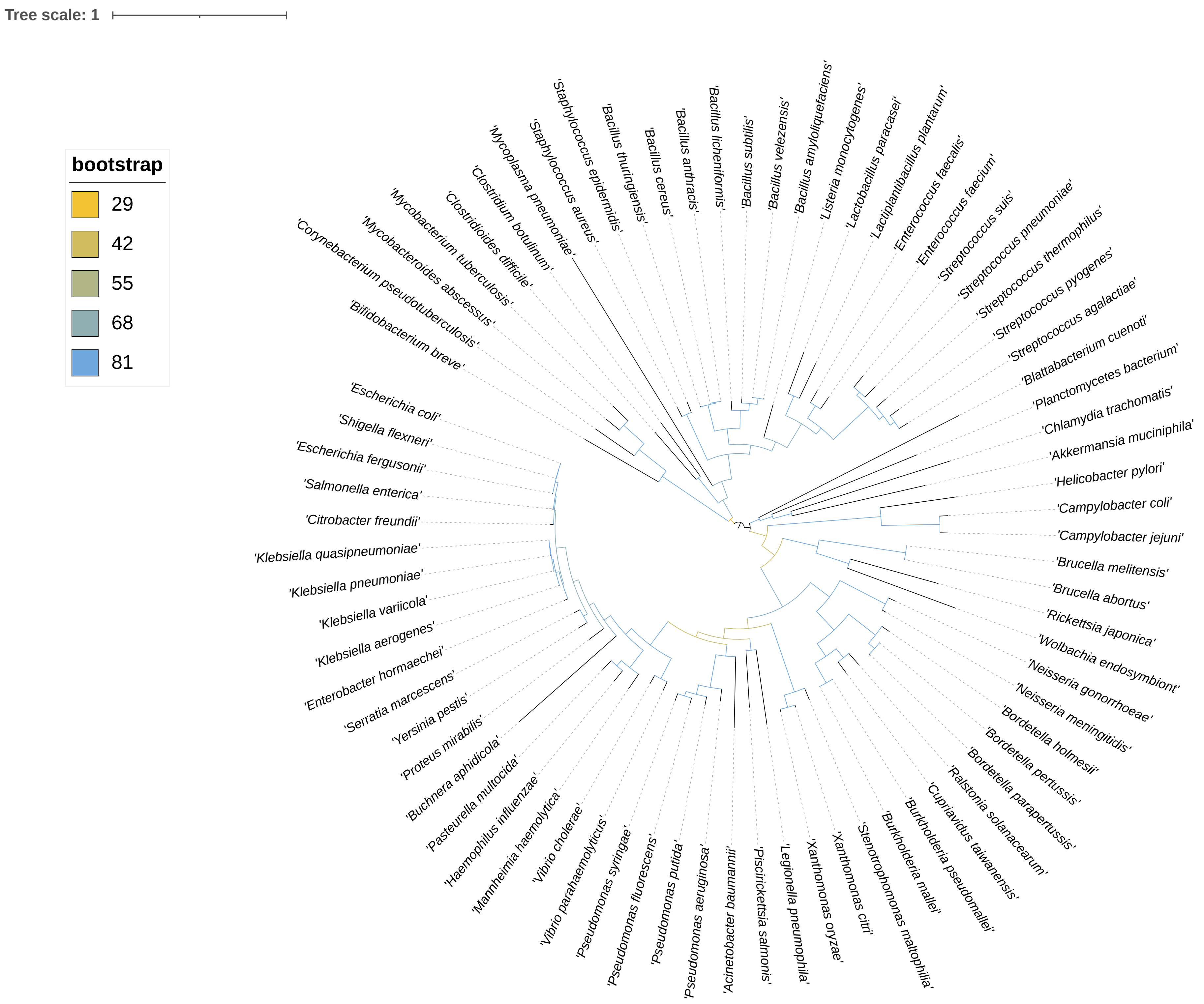
